# Soil prokaryotes associated with decreasing pathogen density during anaerobic soil disinfestation

**DOI:** 10.1101/774810

**Authors:** Chol Gyu Lee, Eriko Kunitomo, Toshiya Iida, Kazuhiro Nakaho, Moriya Ohkuma

**Affiliations:** Graduate School of Bio-Applications and Systems Engineering, Tokyo University of Agriculture and Technology, Koganei, Tokyo, 184-8588, Japan; Japan Collection of Microorganisms, RIKEN BioResource Research Center, Tsukuba, Ibaraki, 305-0074, Japan; Chiba Prefectural Agriculture and Forestry Research Center, Chiba, Chiba 266-0006, Japan; Institute of Vegetable and Floriculture Science, National Agriculture and Food Research Organization, Tsu, Mie 514-2392, Japan

**Keywords:** Bacilli, Clostridia, C/N ratio, Fusarium wilt, rank correlation

## Abstract

Anaerobic soil disinfestation (ASD) is a chemical-independent method that can reduce pathogens. Although soil microbes play essential roles in ASD, the relationship between the microbial community structure and disinfestation efficiency remains unclear. To this end, we investigated changes in the microbial community and pathogen density during a period of ASD under field conditions for 14 days in a greenhouse using three different substrates. Soil samples were collected at 0, 3, 7, and 14 days after ASD treatment. The pathogen densities were analyzed by real-time polymerase chain reactions, prokaryotic community analysis was conducted using unidirectional pyrosequencing, and the factors related to pathogen density were statistically analyzed. The pathogen density rapidly decreased by >90% at 3 days after treatment and then slowly decreased until day 14, but the rate of decrease differed among the substrates. The microbial communities became altered after 3 days and recovered to their original state on day 14. The dipyridyl reaction, microbial diversity, richness, and community structure were not correlated with pathogen density. The most negatively correlated operational taxonomic units with pathogen density were Clostridia and Bacilli, both belonging to Firmicutes. These results suggested that the growth of specific microbes, but not the changes in microbial community structure, might be important for ASD disinfestation efficiency.

## 1. Introduction

Tomato (*Solanum lycopersicum*) is one of the most important vegetables worldwide, with a global yield of approximately 240 million tons in 2017 (FAO, 2019). Soil-borne pathogens cause various plant diseases, including take-all, damping-off, crown rots, and wilting. Fusarium wilt, caused by *Fusarium oxysporum* f. sp. *lycopersici* (FOL) is one of the most serious soil-borne tomato diseases (Larkin and Fravel, 1998). The control methods for this disease, such as soil amendments, crop rotation, biological control, and field sterilization, are often ineffective because they can be affected by environmental factors, such as temperature, precipitation, soil properties, etc. (Campbell, 1994). Although soil disinfestation using chemicals can decrease the pathogen, food safety requirements and the need to reduce environmental pollution have made the development of eco-friendly disinfestation methods crucial (Griffiths et al., 2000; Zhou et al., 2019).

Since 2000, the use of anaerobic soil disinfestation (ASD) to generate anaerobic conditions in soil has been studied in Japan (Momma, 2008; Shinmura, 2000), the Netherlands (Blok et al., 2000; Messiha et al., 2007), and the USA (Butler et al., 2012; Rosskopf et al., 2014; Shennan et al., 2014). ASD involves treating the soil with labile organic carbon, irrigation to saturation using water, and covering with polyethylene mulch film for 2–5 weeks. The organic matter increases microbial respiration, and irrigation purges soil air, while the polyethylene film prevents an inflow of oxygen from the atmosphere, which collectively induce reductive soil conditions. This technique is effective for suppressing several soil-borne diseases, including bacterial wilt, Fusarium wilt, and root rot nematode (Butler et al., 2014; Shrestha et al., 2016). However, the exact mechanisms of ASD that lead to disease suppression remain unclear.

Soil microbes play an essential role in disinfestation by ASD. Momma et al. (2010) showed that sterilized soil loses its disinfestation ability. Microbial communities drastically changed following ASD treatment, especially Clostridia, a class of anaerobic bacteria, that increased significantly after ASD treatment (Mowlick et al., 2013a, 2013b; Rosskopf et al., 2014). However, Clostridia often increase in anaerobic soil regardless of ASD treatment. The relationship between the disinfestation efficiency and microbial community during ASD treatment is unelucidated. To clarify this relationship, monitoring of the changing microbial communities and pathogen densities during the disinfestation period are required. To our knowledge, the changes in the microbial community during the course of ASD treatment have only been analyzed by Li et al. (2017) under *in vitro* conditions, and the relationship between prokaryotic communities and pathogen density was not established.

Ethanol and molasses (as labile organic substrates) and wheat residue, rice husk, and mustard residues (as recalcitrant organic substrates) are often used for ASD, and the substrate type has been found to affect the disinfestation efficiency (Strauss and Kluepfel, 2015; Testen and Miller, 2018). In our study, two labile organic substrates (sugar-contained diatoms [SCDs] and dried molasses [DM]) and one recalcitrant organic substrate (wheat bran [WB]) were used as the carbon sources. We aimed to demonstrate 1) the behavior of FOL and microbial communities over time, and 2) the effects of substrate types on FOL density and microbial community structure during the ASD period under field conditions.

## 2. Materials and methods

### 2.1 Sampling field and ASD treatment

The experiments were performed with soil samples from a tomato-planted greenhouse located in Chiba Prefectural Agriculture and Forestry Research Center, Chiba prefecture (35° 54’ N, 140° 19’ E). The soil pH was 5.9, and the carbon and nitrogen concentrations were 47.2 and 3.38 g kg soil^−1^, respectively. To make FOL-contaminated soil, 10 FOL-infected tomato tubers were packed in a mesh bag and buried in the soil at a depth of 0–15 cm. WB, SCDs (Ajinomoto Coo., Inc., Saga, Japan), and DM (Omalass 95; Westway Feed Products, Tomball, TX, USA) were used as substrates for the ASD treatment. SCDs are discharged from food-processing facilities as by-products of the filtration of saccharified liquids. The main components of such by-products are sugars derived from the saccharified solution of tapioca starch and diatoms used as a filtering aid. These by-products, containing 40% sugar by weight, were powdered. DM is a livestock feed containing water-soluble sugar. The material comprises 33% soybean husks and 67% sugarcane molasses, and they were absorbed. The chemical properties of each substrate are shown in Table 1. The treatment was applied at a rate of 15 t ha^−1^ in each field with a rototiller at a depth of 30 cm. Each treatment was performed in duplicate on 2.5 × 2.2 m plots that were separated by a 50-cm high wave barrier. The field was covered by a 0.1-mm thick transparent polyethylene film and flooded with 150 L of water on day 0. Each site was flooded at the time of disinfestation, and no irrigation was conducted thereafter. Disinfestation was conducted from August 3, 2015, for 14 days. Soil samples were collected at a depth of 0–15 cm from each treated field using a core sampler (Gauge Auger DIK-106B; Daiki Rika Kogyo Co., Ltd, Saitama, Japan) on day 0, 3, 7, and 14 after the ASD treatment. The reduction area was visualized by spraying a bipyridyl solution (1.0 g of 2,2′-bipyridyl and 77 g of ammonium acetate dissolved in 1 L of 1% acetic acid) on a freshly exposed soil face. Soil samples were collected from five randomly selected locations in each plot and mixed well to make a composite sample. A total of 24 soil samples (three substrates × four sampling periods × two replicates) were collected and stored at −20 °C until use.

**Table 1.** Chemical properties of each substrate used for the ASD treatment

| Organic matter | TC<br>(g kg<sup>-1</sup>) | TN<br>(g kg<sup>-1</sup>) | C/N ratio<br>(g kg<sup>-1</sup>) | WSOC<br>(g kg<sup>-1</sup>) |
| --- | --- | --- | --- | --- |
| WB | 400 | 24.0 | 16.4 | 132 |
| SCD | 250 | 10.3 | 24.2 | 181 |
| DM | 419 | 13.8 | 30.4 | 357 |

TC: total carbon, TN: total nitrogen, C/N ratio: the ratio of total carbon per total nitrogen, WSOC: water-soluble organic carbon, WB: wheat bran, SCD: sugar-contained diatoms, DM: dried molasses.

### 2.2 Quantification of *F. oxysporum* in the field

Soil DNA was extracted from 0.5 g of soil with an ISOIL for Beads Beating kit (Nippon Gene, Tokyo, Japan) following the manufacturer’s instructions. DNA quantification and integrity were measured using a Nanodrop spectrophotometer (Thermo Fisher Scientific, Waltham, MA, USA) and gel visualization (0.8% agarose in tris-acetate-EDTA buffer), respectively. Real-time polymerase chain reactions (PCRs) were performed for quantification of *F. oxysporum* density in soil, according to (Inami et al., 2010). The reaction mixture (20 μl) contained 10 μl of TaqMan Universal Master Mix (Thermo Fisher Scientific), 0.5 μM of each primer, 0.25 nM of TaqMan probe, and 15 ng of template DNA. Real-time PCR was performed in duplicate for each sample to amplify the rDNA intergenic spacer region of *F. oxysporum* f. sp. *lycopersici* using specific primer sets SIX1f (5’-GTGCCAGCMGCCGCGGTAA-3’) and SIXir (5’-GGAC-TACVSGGGTATCTAA-3’) and a TaqMan probe carrying a reporter (6-carboxyfluorescein) and a quencher (minor groove binder) SIX1pr (5’-TTGACCTACACGGAATAT-3’). Standard curves were obtained by serial dilutions of linearized plasmids with cloned fragments of the specific genes and were linear (R^2^ = 0.99) in the range used (data not shown).

### 2.3 Tag-encoded amplicon sequencing targeted by 16S rRNA for prokaryotes

PCR was performed on each sample to amplify the V4 variable region of the 16S rRNA gene using the bacterial and archaeal universal primers 515F (5’-GTGCCAGCMGCCGCGGTAA-3’) and 806R (5’-GGAC-TACVSGGGTATCTAA-3’) (Caporaso et al., 2011). Each PCR amplicon was purified twice using the Agencourt AMPure XP system (Beckman Coulter, Inc., Brea, CA, USA) to remove short DNA fragments and was quantified using a Qubit Fluorometer (Invitrogen, Carlsbad, CA, USA). Following successful amplification, the PCR products were adjusted to equimolar concentrations and subjected to unidirectional pyrosequencing at Bioengineering Lab. Co., Ltd. (Kanagawa, Japan) on a MiSeq instrument (Illumina, San Diego, CA, USA). A total of 1,161,378 sequences were obtained from the 24 samples after sequencing (Supplemental Table 1). The sequencing data were deposited in the DNA Data Base of Japan Sequence Read Archive under accession number DRA006673.

### 2.4 Data analysis

Raw FASTQ files were pre-processed using Quantitative Insights Into Microbial Ecology (QIIME) (Caporaso et al., 2010). Data from read sequences, quality, flows, and ancillary metadata were analyzed using the QIIME pipeline. Quality filtering consisted of discarding reads <200 bp or >1000 bp in length, excluding homopolymer runs of >six bp and >six continuous ambiguous bases, and accepting one barcode correction and two primer mismatches. Moreover, reads with a mean quality score <25 were also removed. Finally, singleton operational taxonomic units (OTUs) and chimeric sequences were removed for statistical analysis. Denoising was performed using the built-in Denoiser algorithm, and chimera removal and OTU picking were accomplished with Usearch61 considering a pairwise identity percentage of 0.97. Taxonomy assignment was performed using the Ribosomal Database Project Classifier, a naïve Bayesian classifier, with a minimum confidence of 0.8 against the Greengenes database, October 2012 release. The OTU-based analysis was performed on pyrotag-based datasets to calculate richness and diversity using the phyloseq package of R 3.5.1 (McMurdie and Holmes, 2013). The diversity within each sample was estimated using the non-parametric Shannons’s diversity index and Simpson’s diversity index. The Chao1 estimator was calculated to estimate the species richness of each sample.

Multivariate analysis of community structure and diversity was performed on the pyrotag-based datasets using a weighted UniFrac dissimilarity matrix calculated in QIIME, jackknifing (1000 reiterations) read abundance data at the deepest level possible (9601 reads), and unconstrained ordination offered by cluster analysis with Ward’s method. The effect of differences between substrates and the disinfestation period on the microbial community were assessed by permutational multivariate analysis of variance using R. The Mantel test was used to evaluate the effect of FOL density on microbial communities (Anderson and Walsh, 2013). Spearman’s rank correlation methods were used to determine which OTUs were correlated with FOL density changes common among each substrate-treated soil sample.

## 3. Results

### 3.1 Changes of FOL density during ASD

We described each sample with the substrate name and replication number, for example, WB1 indicated the first sample of two replicates in soil receiving WB treatment. The FOL density was 642–1832 copies g soil^−1^ on day 0, decreasing to 0.6%–10.6% on day 3 and then gradually decreasing up to day 7, except for sample DM1 (Table 2). On day 14, the FOL densities of the DM-treated samples were decreased (<1 copy g soil^−1^), but that of the other treatments were not. A bipyridyl reaction was observed in WB1, SCD1, and DM1 14 days after ASD (Supplemental Table 2).

**Table 2.**
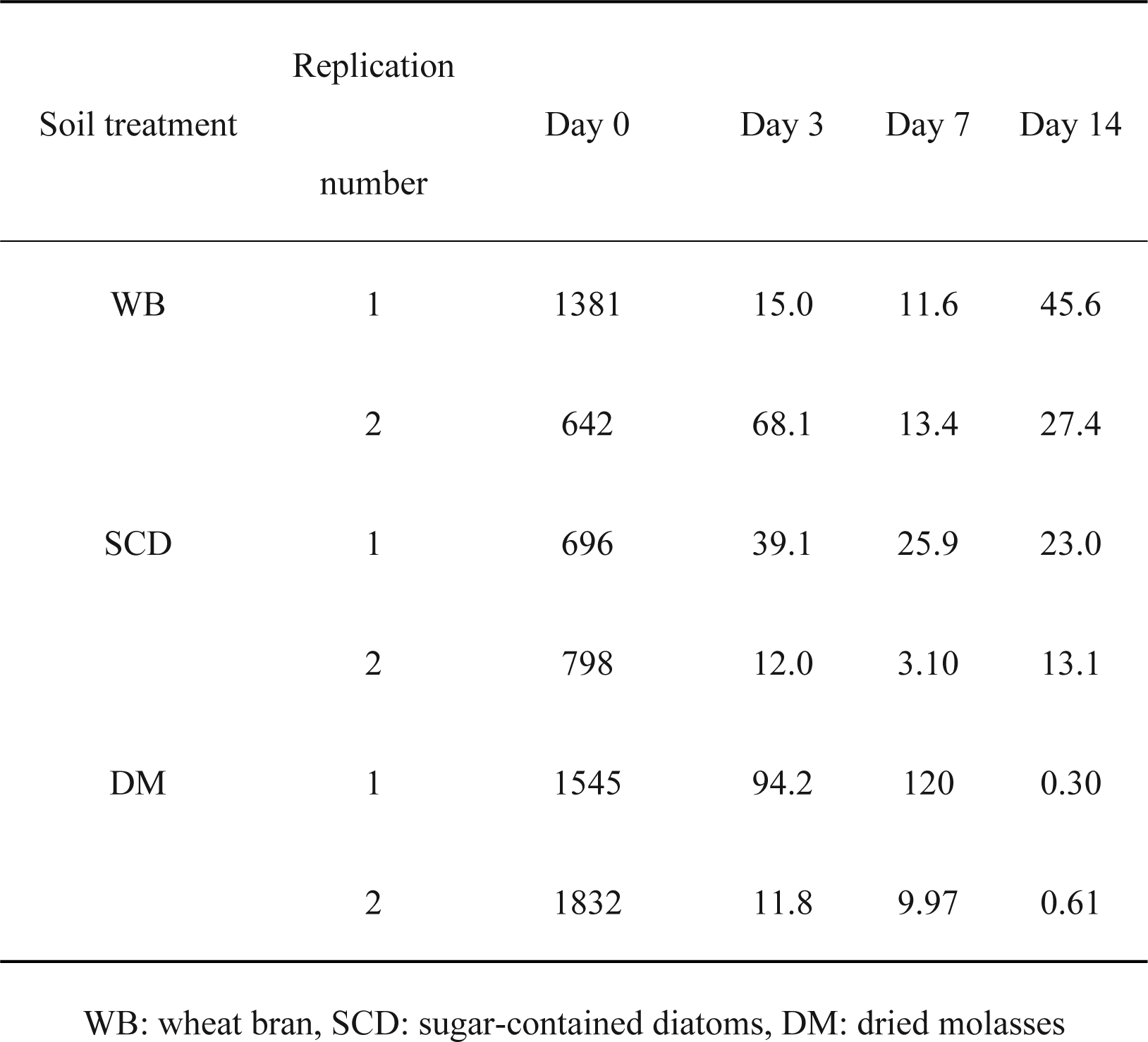
Changes of F. oxysporum density (copies g soil^-1^) during disinfestation period

### 3.2 The succession of prokaryotic soil communities during the disinfestation period

The prokaryotic sequences were clustered into 130,331 OTUs at a 97% similarity. Shannon’s index changed from 0.78 to 1.1 times when day 0 was compared with day 14 (after disinfestation) across each treatment. For the SCD-treated samples, a decrease in both OTU numbers (SCD1, 76% decrease; SCD 2, 32% decrease) and Chao1 (SCD1, 89% decrease; SCD2, 44% decrease) was observed on day 14 when compared with day 0 (Table 3). For the other treatments, the OTU numbers and Chao1 were mildly decreased or increased after disinfestation. Our results also indicated that changes in prokaryotic diversity and richness were not correlated with changes in FOL density during the ASD period (Table 4).

**Table 3.**
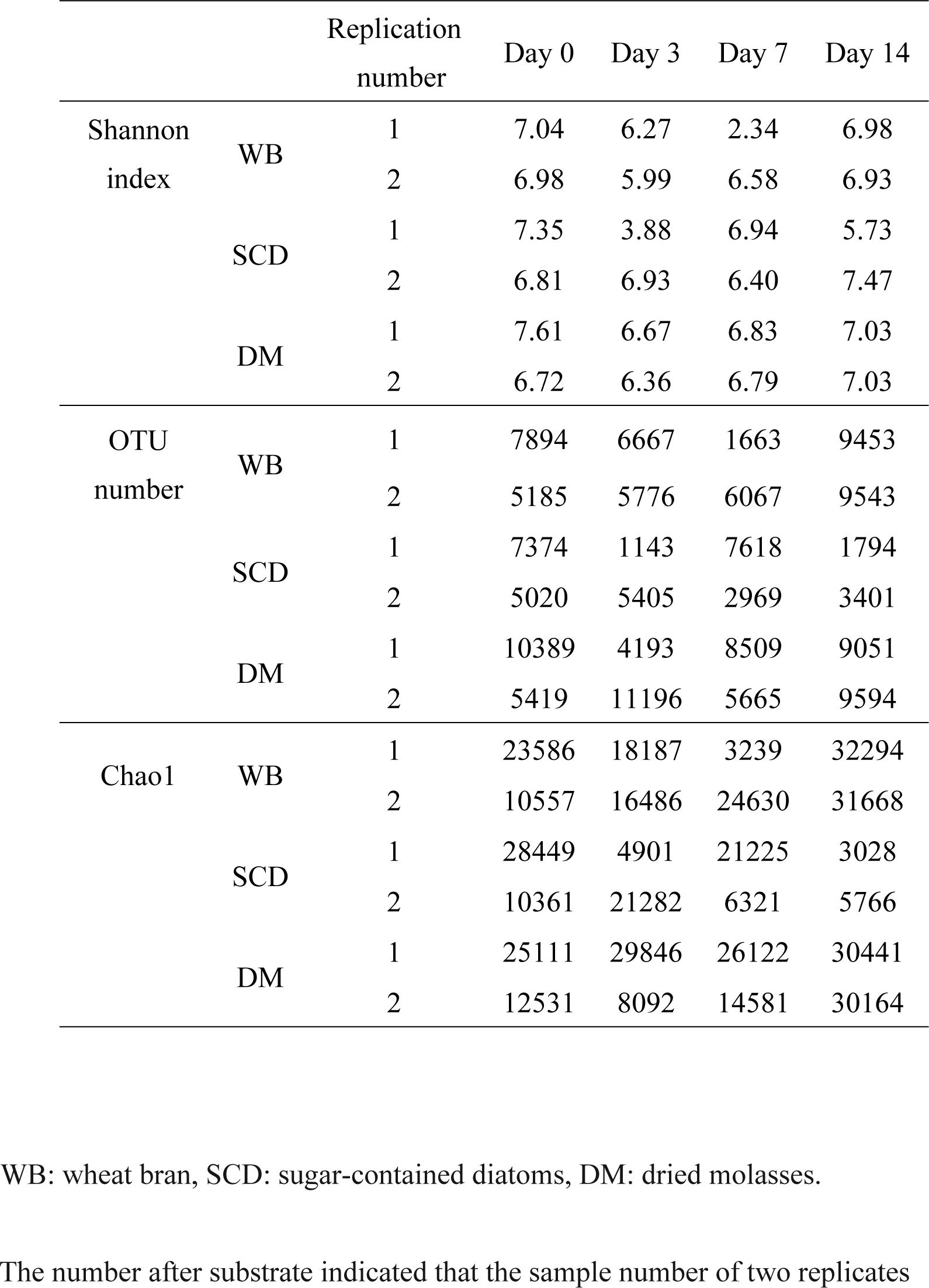
Changes of prokaryotic diversity and richness during the disinfestation period

**Table 4.**
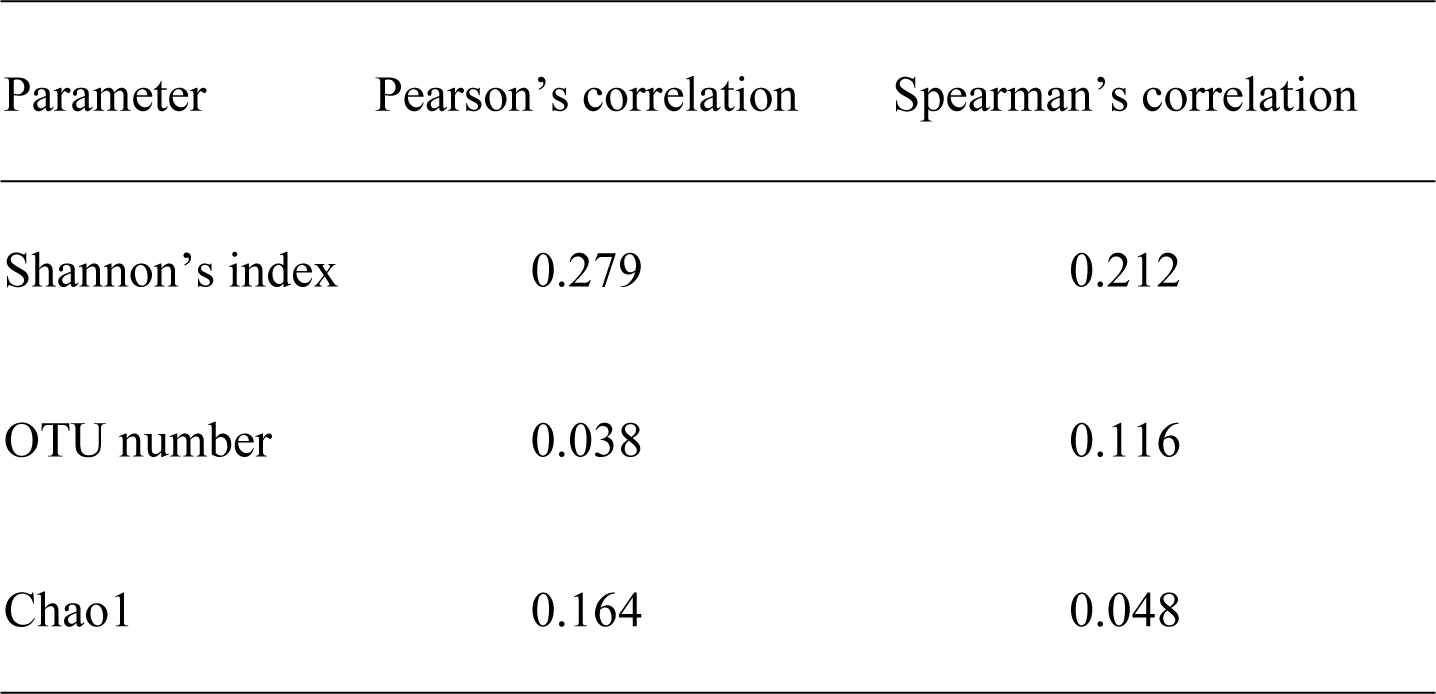
Pearson’s and Spearman’s correlation coefficients between FOL density and prokaryotic diversity or richness

The dominant classes of Bacilli, Clostridia, Alphaproteobacteria, Betaproteobacteria, Deltaproteobacteria, and Gammaproteobacteria occupied >5% of relative abundance in all fields (Figure 1). Bacilli and Clostridia were increased >1.5 times on day 3 in all treatments. In the WB treatment, several microbes were drastically decreased between days 3–7, and they increased from days 7–14. On the other hand, the microbial communities were relatively stable 7 days after the SCD and DM treatments (Figure 1).

**Figure 1.**
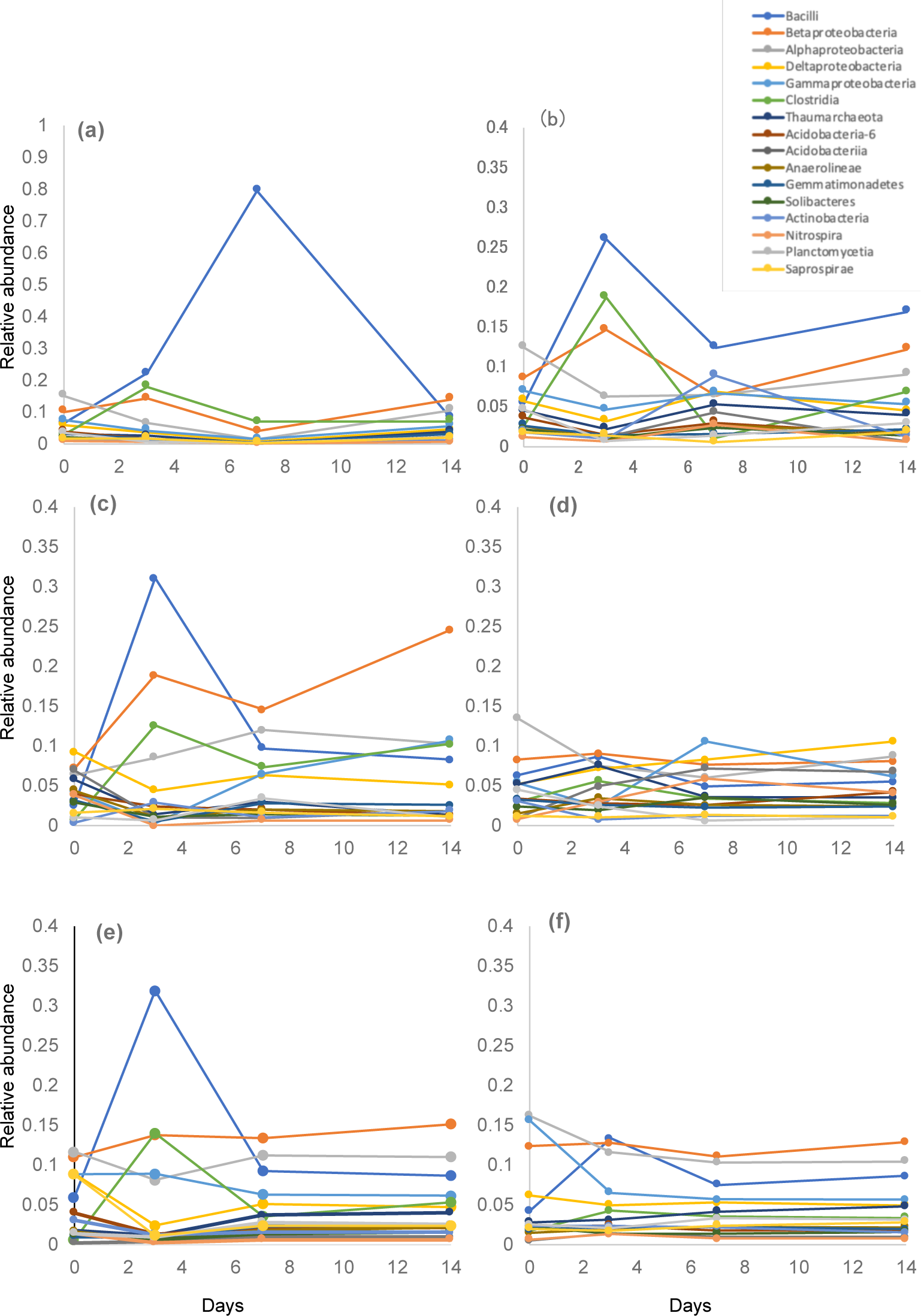
Relative abundance of prokaryotic communities at the class level during the disinfestation period. (a) WB1, (b) WB2, (c) SCD1, (d) SCD2, (e) DM1, (f) DM2

Prokaryotic growth stages were classified as either early (0–3 days), intermediate (3–7 days) or late (7–14 days), based on the period when each prokaryote increased >two-fold for both duplicates. In the SCD and DM treatments, Bacilli and Clostridia were increased in the early stage, and Planctomycetia were increased in the intermediate stage (Table 5). In the WB-treated samples, Bacilli and Clostridia were also increased in the early stage, while Betaproteobacteria, Alphaproteobacteria, Gammaproteobacteria, Planctomycetia, and Saprospirae were increased in the late stage.

**Table 5.** Responses of bacterial classes during ASD

| Bacterial class | WB | SCD | DM |
| --- | --- | --- | --- |
| Bacilli | Early | Early | Early |
| Clostridia | Early | Early | Early |
| Betaproteobacteria | Late | Early |  |
| Alphaproteobacteria | Late |  |  |
| Gammaproteobacteria | Late | Intermediate |  |
| Planctomycetia | Late | Intermediate | Intermediate |
| Saprospirae | Late |  |  |

The bacteria that increased >2-fold times in both duplicates were selected.
Early: day 0–3; intermediate: day 3–7 day; late: day 7–14.

Weighted UniFrac analysis showed that prokaryotic communities were roughly separated throughout the disinfestation period (Figure 2). The initial microbial community was similar in all samples except for SCD1 and became separated as SCD1, WB1, WB2, and DM2 on day 3, as was the case on day 0. The changes in the prokaryotic community in the late stage were not different in DM1, DM2, and SCD2 because they were included in the same cluster on day 7 and 14. On the other hand, that of the other treatments were divided into different clusters between day 7 and 14.

**Figure 2.**
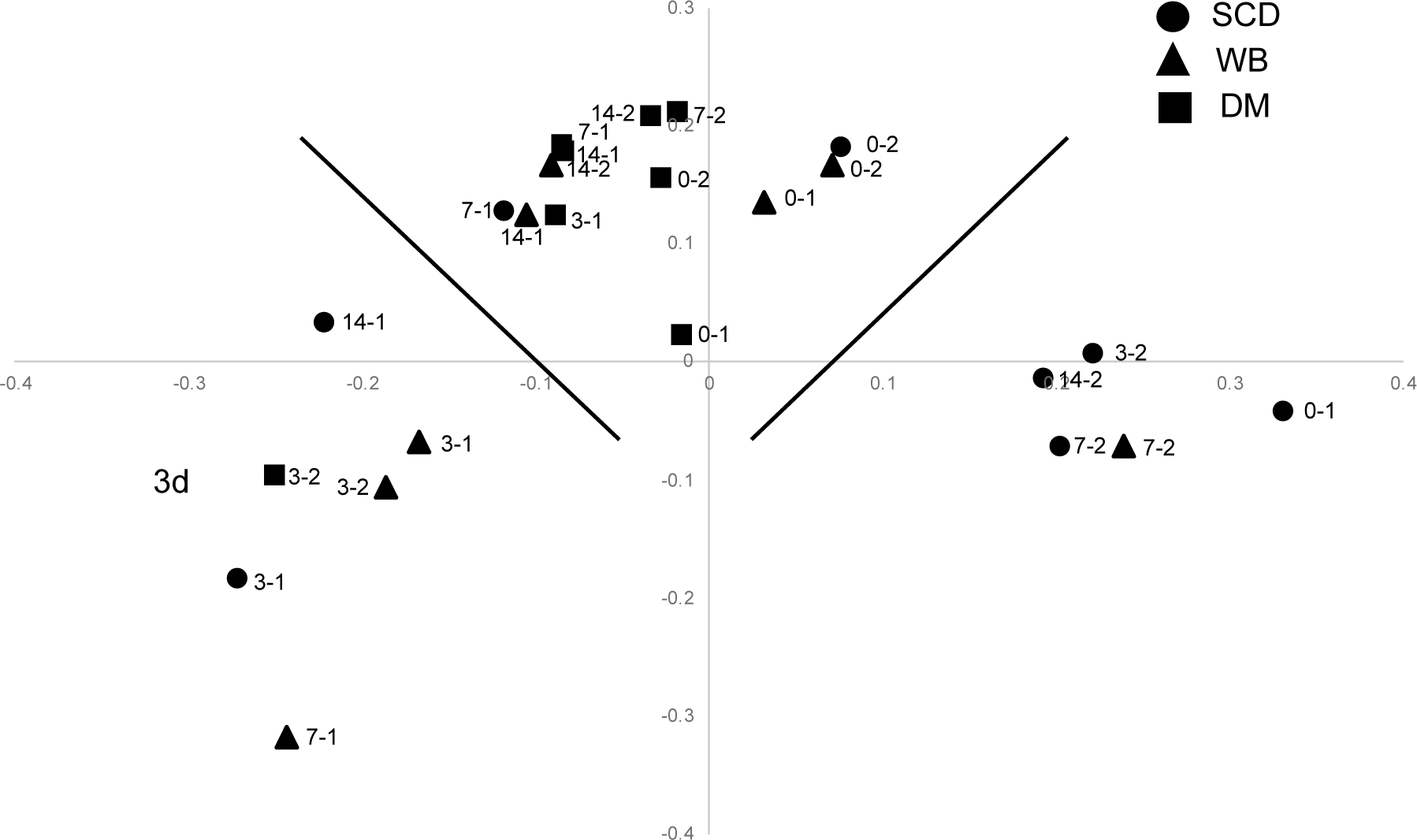
Changes in prokaryotic communities during ASD treatment. WB: wheat bran, SCD: sugar-contained diatoms, DM: dried molasses. The numbers at each plot show the date of sampling and the sample number connected with a hyphen. The line indicates the separation of each cluster based on *k*-means analysis.

PERMANOVA analysis revealed that the microbial communities were independently affected by substrates and the disinfestation period (Table 6). The Mantel test showed that the microbial communities were not altered by FOL density (F = 1.82, p = 0.06). These results suggested that factors other than the microbial community affected the decrease in FOL density. Then, OTUs that changed along with FOL density were selected using rank correlation analysis. Eleven OTUs were negatively (*ρ* < −0.5) correlated with changes in FOL density that were common among each substrate-treated field (Table 7). There were 7/11 OTUs belonging to Firmicutes; of which three (19544, 122066, and 95490) belonged to Clostridia, 2 (7428 and 109424) to Bacilli, and 1 (78089) to Negativicutes. The other OTUs belonged to Acidobacteria (24789), Planctomycetia (9173), and Proteobacteria (69473 and 52717).

**Table 6.**
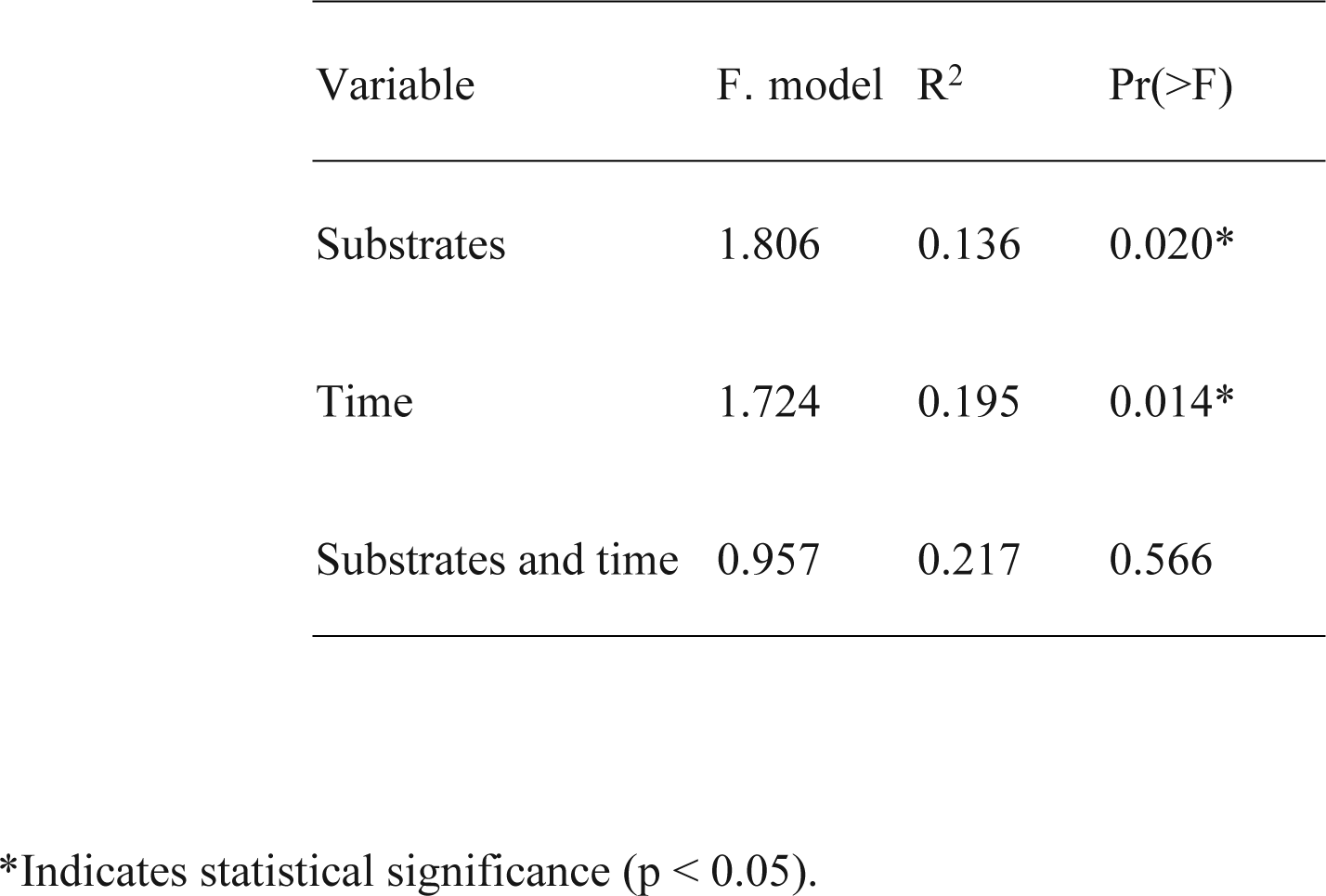
PERMANOVA results for prokaryotic communities

**Table 7.** OTUs negatively correlated with FOL density

| OTU number | Rho value | Closest relatives | Phylum |
| --- | --- | --- | --- |
| OTU19544 | −0.585 | Clostridiales | Firmicutes |
| OTU122066 | −0.563 | *Clostridium* | Firmicutes |
| OTU7428 | −0.558 | Bacillales | Firmicutes |
| OTU109424 | −0.558 | *Paenibacillus* | Firmicutes |
| OTU78089 | −0.556 | Veillonellaceae | Firmicutes |
| OTU23993 | −0.536 | *Unidentified Firmicutes* | Firmicutes |
| OTU95490 | −0.528 | Peptococcaceae | Firmicutes |
| OTU24789 | −0.527 | *Candidatus Solibacter* | Acidobacteria |
| OTU9173 | −0.508 | Planctomycetaceae | Planctomycetia |
| OTU69473 | −0.507 | Unidentified<br>Proteobacteria | Proteobacteria |
| OTU52717 | −0.502 | Comamonadaceae | Proteobacteria |

Rho value indicates the correlation coefficient between FOL density and each OTU as per Spearman's rank correlation method.

## 4. Discussion

### 4.1 Relationships between substrate types, microbial community, and FOL density

The C/N ratio of substrates is one of the indicators of the decomposition rate (Constantinides and Fownes, 1994; Nicolardot et al., 2001). The WB-treated samples showed a lower level of water-soluble organic carbon and a higher C/N ratio than those treated with SCD or DM because WB is composed of recalcitrant carbon fractions, such as cellulose, hemicellulose, and lignin. Following WB treatment, the microbial community was changed, even after 7 days, but this was unrelated to the disinfestation effect. During ASD, DM rapidly dissolved in the irrigation water because of a higher water-soluble organic carbon fraction and C/N ratio. Following DM treatment, FOL density decreased during days 7–14, but the structures of the prokaryotic communities were not drastically changed during these periods because of the rapid decomposition of the substrate. Previous studies have shown that organic matter with a lower C/N amended into the soil could induce the highest anaerobic conditions, and the substrate C/N ratio was negatively correlated with the disinfestation effects (Blok et al., 2000; Shrestha et al., 2016; Testen and Miller, 2018). In our study, different substrates exerted different effects on the prokaryotic community, but the substrate decomposition rate, especially the variables of substrate C/N ratio and the content of water-soluble organic carbon, was not correlated with the disinfestation efficiency.

### 4.2 Relationships between microbial diversity, community structure, and FOL density

During ASD treatment, as the microbial community shifts toward facultative and obligate anaerobes, the anaerobic decomposition of labile carbon creates short-chain organic acids (e.g., acetic, n-butyric, and propionic acid). The volatile fatty acids (VFAs) are likely toxic to soil-borne plant pathogens, plant-parasitic nematodes, and weeds (Momma, 2008; Momma et al., 2006). (Momma et al., 2013) showed that bipyridyl testing (an indicator of reduction) of disinfested soil was a useful method to evaluate whether ASD treatment was conducted appropriately. Liu et al. (2016) showed that although Shannon’s diversity index was decreased on day 4 and did not change significantly thereafter, the FOL population decreased after day 4. Messiha et al. (2007) indicated that microbial diversity was not different between substrate amended and non-amended soil following ASD treatment, but their community structure was different. On the other hand, Shennan et al. (2014) showed that ASD conducted with molasses did not alter community structure. These studies suggest that microbial diversity, community structure, and soil redox potential (Eh) are not correlated with disease incidence, corresponding with our findings. Hence, different mechanisms could be critical for suppressing specific organisms, but the production of organic acids via the anaerobic decomposition of added carbon, release of VFAs, and biocontrol by microorganisms that flourish during the process are all potentially important (Momma et al., 2007; Shrestha et al., 2016). Yonemoto et al. (2006) indicated that a decreasing Eh value and FOL density were not directly correlated and concluded that it was important for the concentration of VFAs or microbial community change. Li et al. (2017) revealed the changes in the microbial community during the disinfestation period, and the presence of some anaerobic bacteria correlated with soil organic acid content.

Unfortunately, we did not analyze the soil chemical properties, such as pH, VFA concentrations, and soil organic carbon concentration, in our study. Recent studies have shown that specific microbes, like Clostridia and *Zopfiella*, isolated from ASD-treated soil could suppress disease incidence (Liu et al., 2019; Momma, 2008; Ueki et al., 2017). Therefore, specific microbes may be necessary for disease suppression by ASD.

### 4.3 Microbes associated with FOL dynamics

Studies have shown that Clostridia and Bacilli belonging to Firmicutes increased in number and became the dominant bacteria following ASD treatment regardless of soil and substrate types (Huang et al., 2016, 2015; Mowlick et al., 2012; Poret-Peterson et al., 2019, Testen and Miller, 2018). During ASD, decreased oxygen promotes the increased prevalence of anaerobic microbes (Momma et al., 2006; Mowlick et al., 2014, 2013a; Runia et al., 2014). In our study, the conditions induced by ASD, regardless of the carbon source, may have functioned as a habitat filter, allowing the proliferation of closely related taxa with shared physiological adaptations. Clostridia and Bacilli drastically increased and FOL decreased in the early stage of ASD. Increasing clostridial populations in ASD-treated soils might correlate with the elevated production of VFAs and is likely toxic to soil-borne plant pathogens (Momma et al., 2006; Mowlick et al., 2012). Some Clostridia can directly kill FOL (Momma, 2008; Ueki et al., 2017). Bacilli, including *Bacillus* and *Paenibacillus*, are well-known biocontrol agents for FOL. Some Bacilli might contribute to the rapid decrease of soil Eh at the initial stage of ASD treatment via oxygen consumption (Mowlick et al., 2012).

Therefore, increased Clostridia and Bacilli may induce a decrease in FOL density in the early stage of ASD treatment by decreasing oxygen, producing VFA, and directly killing FOL. We have yet to obtain direct evidence for the suppression of FOL by Clostridia and Bacilli, but these biocontrol agents may play essential roles in disinfestation.

## 5. Conclusion

Soil microbes play important roles in ASD efficiency. In this study, we analyzed the changes of FOL density and microbial community during ASD among three substrates to elucidate the relationship between FOL density and the microbial community. FOL density was drastically decreased for the first 3 days following ASD and slowly continued decreasing until day 14. The microbial community was significantly changed on day 3, but some microbes were increased until day 14 following WB treatment. The microbial diversity, richness, and community structure as well as the C/N ratio of substrates were not correlated with FOL density in all treatments. Notably, Clostridia and Bacilli were negatively correlated with a decrease in FOL density. These results suggested that specific microbes might be involved in disinfestation efficiency and not changes in the entire community structure itself. Future studies will investigate the usefulness of these microbes as indicators of ASD efficiency and their roles in disease suppression.

## Acknowledgments

This work was supported by the Cabinet Office, Government of Japan, Cross-ministerial Strategic Innovation Promotion Program (SIP), and the “Technologies for creating next-generation agriculture, forestry and fisheries” (funding agency: Bio-oriented Technology Research Advancement Institution, NARO).

This work was also partially supported by the RIKEN Competitive Program for Creative Science and Technology, Grants-in-Aid for Scientific Research from the Japan Society for Promotion of Science, Nos. 17K01447 and 19H05689 (to M.O.). The part of prokaryotic community analysis was supported by Dr. Kozue Sawada and Ms. Hana Kobayashi at Tokyo University of Agriculture and Technology.

---

**Supplemental Table 1.** Sequencing read numbers of each sample

|  | Replication<br>number | Day 0 | Day 3 | Day 7 | Day 14 |
| --- | --- | --- | --- | --- | --- |
| WB | 1 | 67,737 | 53,661 | 32,042 | 57,352 |
|  | 2 | 47,167 | 45,457 | 53,315 | 53,786 |
| SCD | 1 | 59,144 | 54,281 | 54,852 | 16,094 |
|  | 2 | 44,798 | 45,825 | 52,338 | 9,601 |
| DM | 1 | 61,651 | 24,593 | 38,626 | 50,307 |
|  | 2 | 40,350 | 89,546 | 56,514 | 55,341 |

WB: wheat bran, SCD: sugar-contained diatoms, DM: dried molasses.

**Supplemental Table 2.**
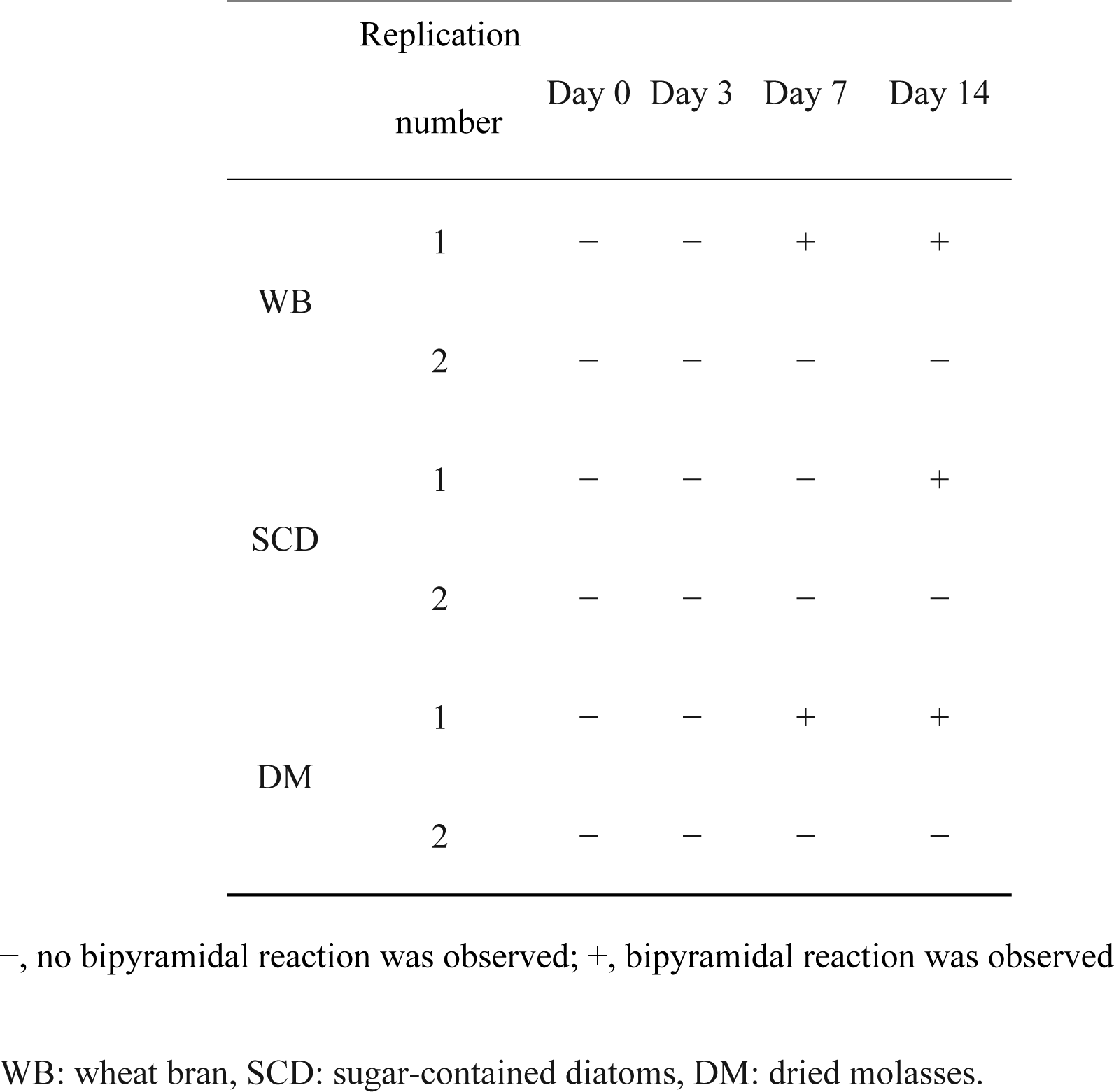
The dipyridyl reaction of each soil sample during disinfestation period

